# Ocular and Facial Far-UVC Doses from Ceiling-Mounted 222 nm Far-UVC Fixtures

**DOI:** 10.1101/2022.05.09.491217

**Authors:** Michael A. Duncan, David Welch, Igor Shuryak, David J. Brenner

**Author notes:** Corresponding Author: David Welch, Center for Radiological Research, Columbia University Irving Medical Center, New York, New York 10032, USA.

## Abstract

Far-UVC radiation, commonly defined as wavelengths from 200 nm – 235 nm, is a promising tool to help prevent the spread of disease. The unique advantage of far-UVC technology over traditional ultraviolet germicidal irradiation lies in the potential for direct application of far-UVC into occupied spaces since antimicrobial doses of far-UVC are significantly below the recommended daily safe exposure limits. This study used a ceiling-mounted far-UVC fixture emitting at 222 nm to directly irradiate an indoor space and then evaluated the doses received upon a manikin. Radiation sensitive film was affixed to the head, nose, lip, and eyes of the manikin, and the 8-hour equivalent exposure dose was determined. Variables examined included manikin height (sitting or standing position), manikin offset from directly below the fixture, tilt of the manikin, the addition of glasses, the addition of hair, and different anatomical feature sizes. Importantly, at the manikin position with the highest dose to eyes, the average eye dose was only 5.8% of the maximum directly measured dose. These results provide the first experimental analysis of possible exposure doses a human would experience from an indoor far-UVC installation.

## Introduction

Preventing the spread of disease is a core public health goal, and the 2019 outbreak of SARS-CoV-2 has highlighted the need for technology to help in this endeavor. Ultraviolet radiation (UV, UVR) is an established method for the inactivation of viruses, bacteria, and other microbes (1). UV wavelengths range from 100 nm to 400 nm, however the 100 nm – 200 nm range is referred to as “vacuum UV” and is not typically included in topics of germicidal UV applications because it is highly absorbed in air. UV wavelengths are commonly categorized as UVC from 200 nm – 280 nm, UVB from 280 nm – 315 nm, and UVA from 315 nm – 400 nm (1). Within the UVC is a subset of wavelengths from 200 nm to 235 nm which is often referred to as the far-UVC. The safety and antimicrobial efficacy of far-UVC wavelengths is an emerging area of research both by our group (2–8) and many others (9–17).

Safety concerns associated with UVR exposure when using UV germicidal lamps are warranted. UV exposure can cause various skin conditions, such as erythema and photoaging, or eye effects, such as photokeratitis, conjunctivitis, cataracts, and pterygium (18–23). Luckily, anatomical features associated with the eye including brow ridge, cheek angel, curvature of the anterior ocular surface, ocular media, ocular protein structure composition, percent of lid closure during exposure, as well as facial coverings such as hats and glasses all play a role in reducing ocular UVR doses (24). While limiting UV exposure to the eyes is generally important for safety, the health hazard for ocular damage is also highly wavelength dependent. Traditional germicidal UV fixtures, usually comprised of low-pressure mercury lamps, emit with a primary peak in the UVC range at 254 nm and are well established for use as physical disinfectants for contaminated air and surfaces(1, 25, 26). However, due to the health hazards of very low doses of 254 nm radiation to the human eye and skin, these UV sources are typically only used to expose unoccupied areas.

Recently, studies of UVR have revealed that 222 nm far-UVC radiation is highly effective for inactivation of surface and airborne microbes while also proving to be much safer to skin and eye tissues. The safety of the far-UVC is derived from the high attenuation of these wavelengths in biological tissues. Radiation at 222 nm can penetrate smaller microbes which are generally less than 1 um in diameter; however, its penetration into human cells (~10-20 um) is minimal because it is largely absorbed by the proteins of the cytoplasm before reaching DNA in the cell nucleus(27–29). This principle of absorption providing protection is already observed more generally for ocular exposures, with the human cornea (approximately 500 μm thick) completely absorbing all UVC radiation so there is not a threat of damage to the lens; far-UVC has an even higher attenuation than other UVC so the penetration of these wavelengths is even more limited (30, 31).

Regardless of wavelength, exposure limits and safety guidelines remain for all UV radiation exposures. Verification that human exposure is kept below these limits is therefore necessary for any installation of UV lighting (32). Occupational safety guidelines, including those published by the American Conference of Governmental Industrial Hygienists (ACGIH)(33) and the International Commission on Non-Ionizing Radiation Protection (ICNIRP)(34), provide widely accepted safe exposure levels for UV lighting. These safe exposure limits are called threshold limit values (TLVs) by the ACGIH and exposure limits (ELs) by the ICNIRP. Prior to this year the TLVs and ELs were generally in agreement for values throughout the UV, but the ACGIH announced higher limits for many UVC wavelengths in their 2022 publication (34–36). The TLV (or EL) specifies the 8-hour exposure limit under which it is expected that humans may be repeatedly exposed without acute effects or risk of delayed effects. The recommendations are wavelength dependent and limits for far-UVC exposure are higher than for conventional germicidal UV; for example, the ICNIRP EL for 222 nm light is 23 mJ/cm^2^ while the EL for 254 nm light is only 6 mJ/cm^2^. Ultimately, the exposure dose received is a product of the source intensity and the exposure duration, so minimizing these factors is often a goal to increase safety. A common use of UVC for germicidal application is upper-air room disinfection. Fixtures for upper room installations are designed to only irradiate and disinfect room air above the occupied space, and this approach effectively limits the dose received by the room occupants. However, because far-UVC installations are being designed to irradiate human occupied spaces, there is a high level of importance to exploring the expected doses an occupant would receive.

Most studies of UV dosing to the human face and eyes have been performed outside utilizing sunlight. A 1988 study by Rosenthal *et al.* examined ocular exposure from UVR wavelengths of 295 - 330 nm on UV sensitive films placed between the eyes of fishermen, landscapers, and construction workers during different times of the year and in the presence and absence of a hat (37). Their study found that wearing a brimmed hat, working in the presence of reflective surfaces, and seasonal variation all significantly affected ocular exposure to UVR. Other more recent studies continue to utilize similar solar principles for ocular exposure, safety, and dosimetry comparisons (38–45).

With the increase in manufacturing of far-UVC emitting lamps, the concept of using 222 nm KrCl excimer lamps in indoor settings for antimicrobial disinfection in the presence of humans is now becoming practical. The installation of far-UVC lamps to directly expose human occupied areas like public transportation, restaurants, and schools appears to be a promising approach to reduce disease transmission. Yet, the UV dose to the human face and eyes from these fixtures mounted in indoor settings remains poorly understood. Here, we use UV sensitive film and a ceiling-mounted 222 nm far-UVC KrCl excimer lamp above realistic human manikin analog heads to determine radiation doses received to the human eyes as well as the head, nose, and lips. The study examines the amount of UV dose received in an extrapolated 8-hour period by the human head at different positions and with varying human features, highlighting the limited dose received by the eyes. Our results provide novel insight regarding safe operation of 222 nm wavelength far-UVC lamps in human occupied spaces.

## Materials and Methods

### UV source

A Lumenizer-300 excimer lamp fixture (LumenLabs, Austin, TX) was used as the UV source for all experiments. The fixture is 305 mm x 345 mm and contains 3 LumenLabs Lpack-1A KrCl excimer bulbs emitting primarily at 222 nm as well as integrated optical filters to reduce emissions outside the 222 nm peak. The 3 lamps are capable of directional adjustments to tilt from a directed downward position to a 45-degree outward position. All experiments were performed with the lamps tilted at the 45-degree outward position. We considered directly under the lamp to be the central axis of the lamp. Lamps were operated for a 20-minute warm-up period before any measurements.

### Light characterization

Spectral characterization of the Lumenizer-300 lamp was performed using a Gigahertz Optik BTS2048-UV spectroradiometer (Gigahertz-Optik Inc, Amesbury, MA). The spectral output for the lamp used in the study has a peak emission at 222 nm, and the UV spectrum is shown in the Supplemental Materials Figure S1. A UIT2400 meter (Ushio America Inc., Cypress, CA, USA) equipped with an SED220 detector and W diffuser input optic was also used for irradiance measurements. The UIT2400 meter was used to obtain horizontal irradiance (meter oriented towards ceiling) and vertical irradiance (meter towards wall) measurements at each of the six measurement positions shown in Figure 1, as well as the direct irradiance measurement by orienting the meter directly at the fixture.

**Figure 1.**
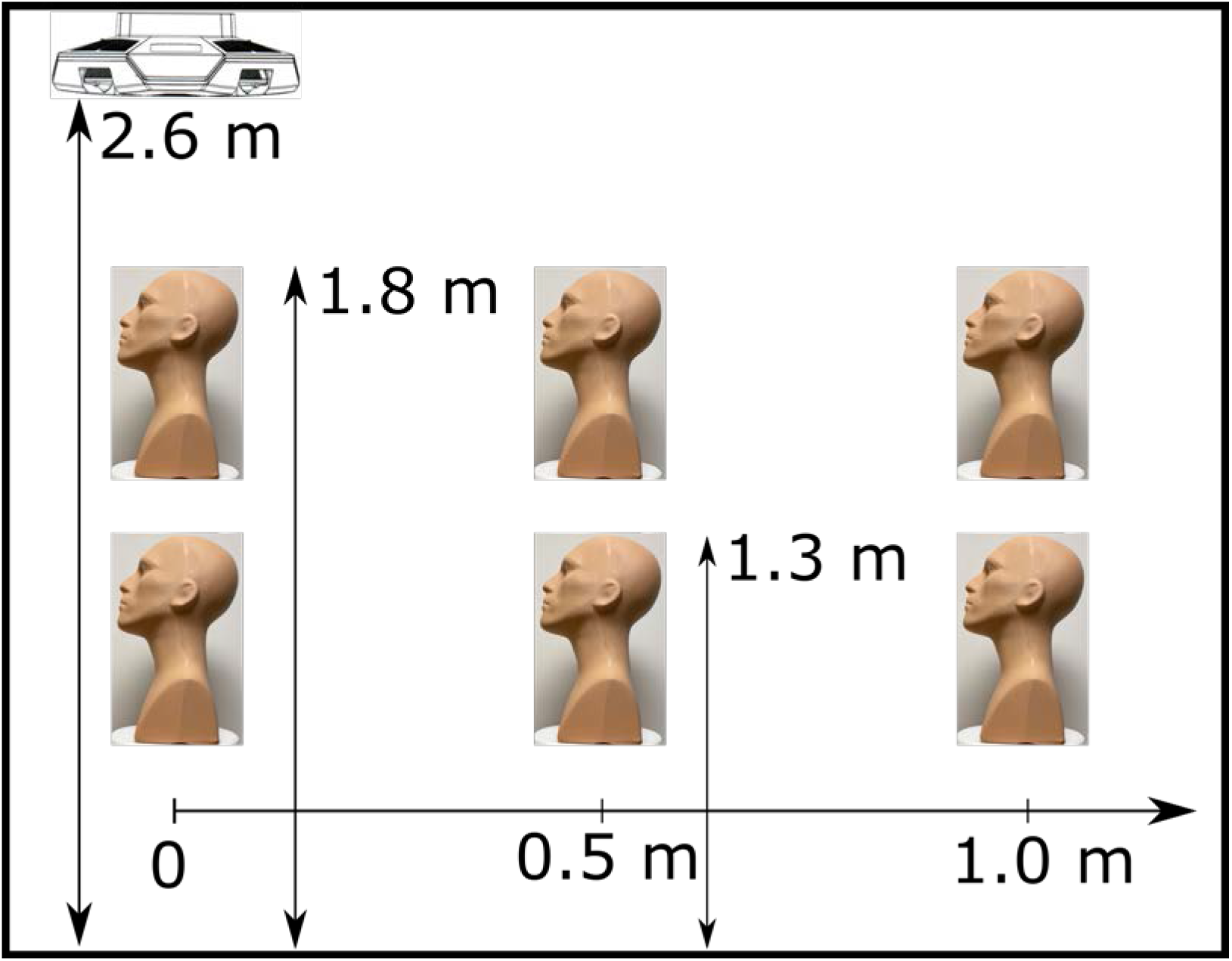
The positions of the far-UVC fixture and the manikin. The far-UVC fixture was attached to the ceiling and directed downward. The six positions of the manikin represent sitting height (1.3 m) and standing height (1.8 m) measured from the floor of the room for three horizontal positions ranging from directly under the lamp to a 1.0 m offset.

### Measurements, location, and positioning

Two manikins with realistic facial features were purchased and used as human analogs for dosing measurements: a primary, bald manikin (Figure 2, panels A-E), and a secondary manikin with in-built hair as seen in Figure 2F. For each manikin, the top of the head was placed at six positions: three positions at approximately the 75^th^ percentile average standing height of adult males (~1.8-meters from the floor) and three positions at a sitting height (~1.3-meters from the floor) (46). Both heights had three different offsets from underneath the central axis of the lamp: 0-meters (directly under the central axis of the lamp), a 0.5-meter offset, and a 1-meter offset. For each position, the manikin was placed on a 360-degree rotating stage which operated continuously at approximately 2 rotations per minute during exposures. The schematic of the positions is provided in Figure 1. Additionally, each location and position had three different tilt angles: a 0-degree tilt, a 15-degree forward tilt, and a 15-degree backward tilt as seen in Figure 2, panels A-C. In all, this totaled 18 unique positional measurements. All experiments used a film exposure time of 10 minutes.

**Figure 2.**
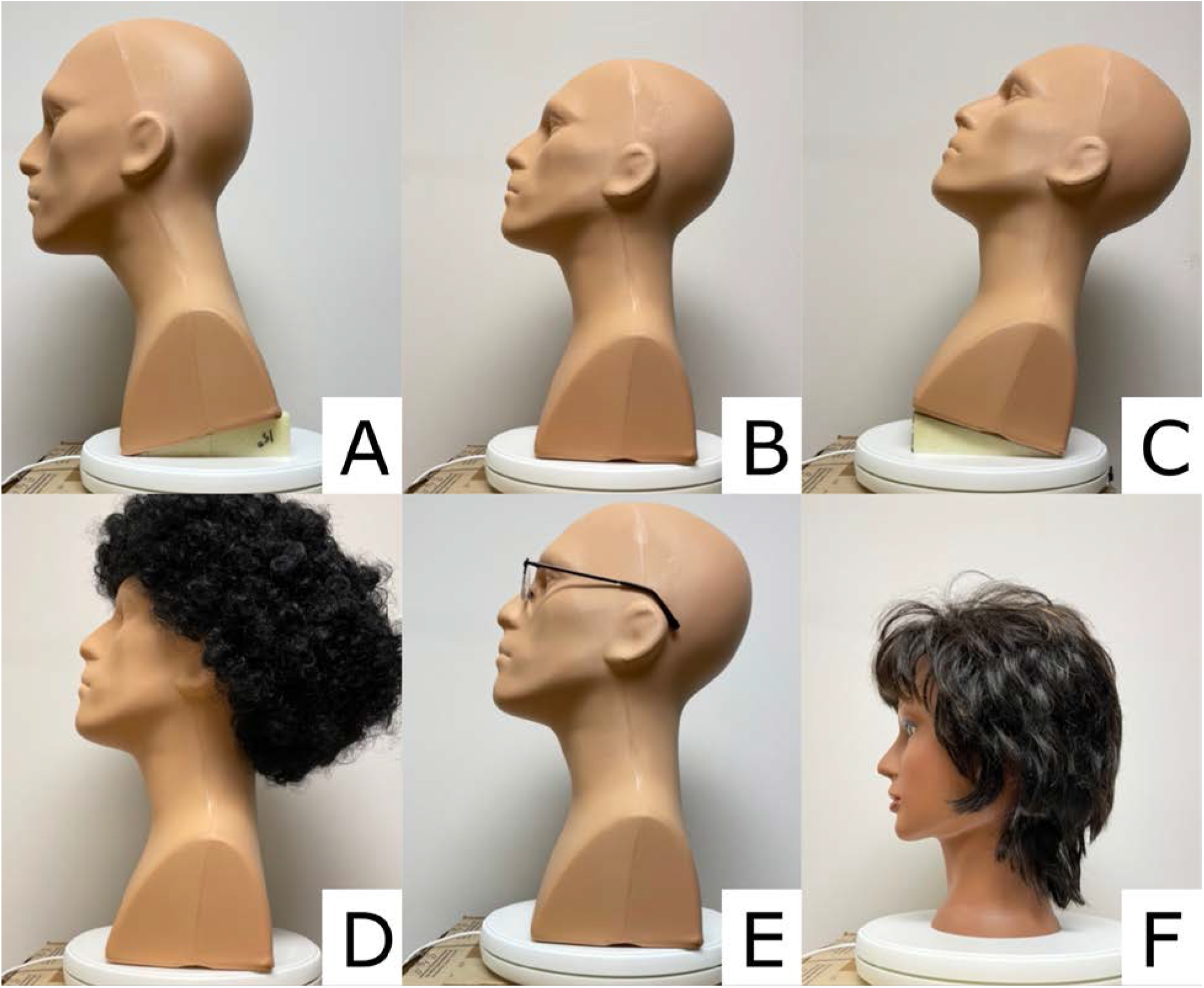
Variations in the manikin setup positions. The top row shows the primary manikin tilted forward 15° (A), not tilted (B), and tilted backwards 15° (C). The bottom row shows the addition of the afro wig to the primary manikin (D), the addition of a pair of glasses to the primary manikin (E), and the secondary manikin with in-built hair (F).

Potential alteration of the UV radiant exposure dosing to the manikin face was also tested with the addition of glasses and different hair style conditions separately. A basic pair of plastic, drug-store reading glasses was used as seen in Figure 2, panel E. One hair style tested was a curly, afro-like wig placed on the head of one primary manikin (Figure 2, panel D); the other hair style was short hair permanently attached to the secondary manikin (Figure 2, panel F). Each additional condition was measured as before for radiant exposure with film in each of the 18 positional locations as mentioned above.

### UV sensitive films

The film product used in this study is OrthoChromic Film OC-1 (Orthochrome Inc., Hillsborough, NJ). OC-1 film is marketed as a tool for radiation therapy measurements though its utility in UVC dosimetry has been demonstrated elsewhere (32). The flexible film is 155 μm thick, consisting of a 30 μm active coating on a 125 μm white polyester base. The active region must be oriented towards the UV source during measurements since the polyester layer is opaque.

Films were cut into rectangles approximately 2 cm x 1.5 cm and placed at the same five anatomical locations on the manikin for each measurement location and position: top of the head, over the left eye, over the right eye, bridge of the nose, and superior lateral position of the upper lip as shown in Figure 3. New films were used for each measurement, and film placement was performed in the absence of lamp radiation to avoid unwanted exposure to active regions before recorded measurements. Films were analyzed within 24 hours of exposure.

**Figure 3.**
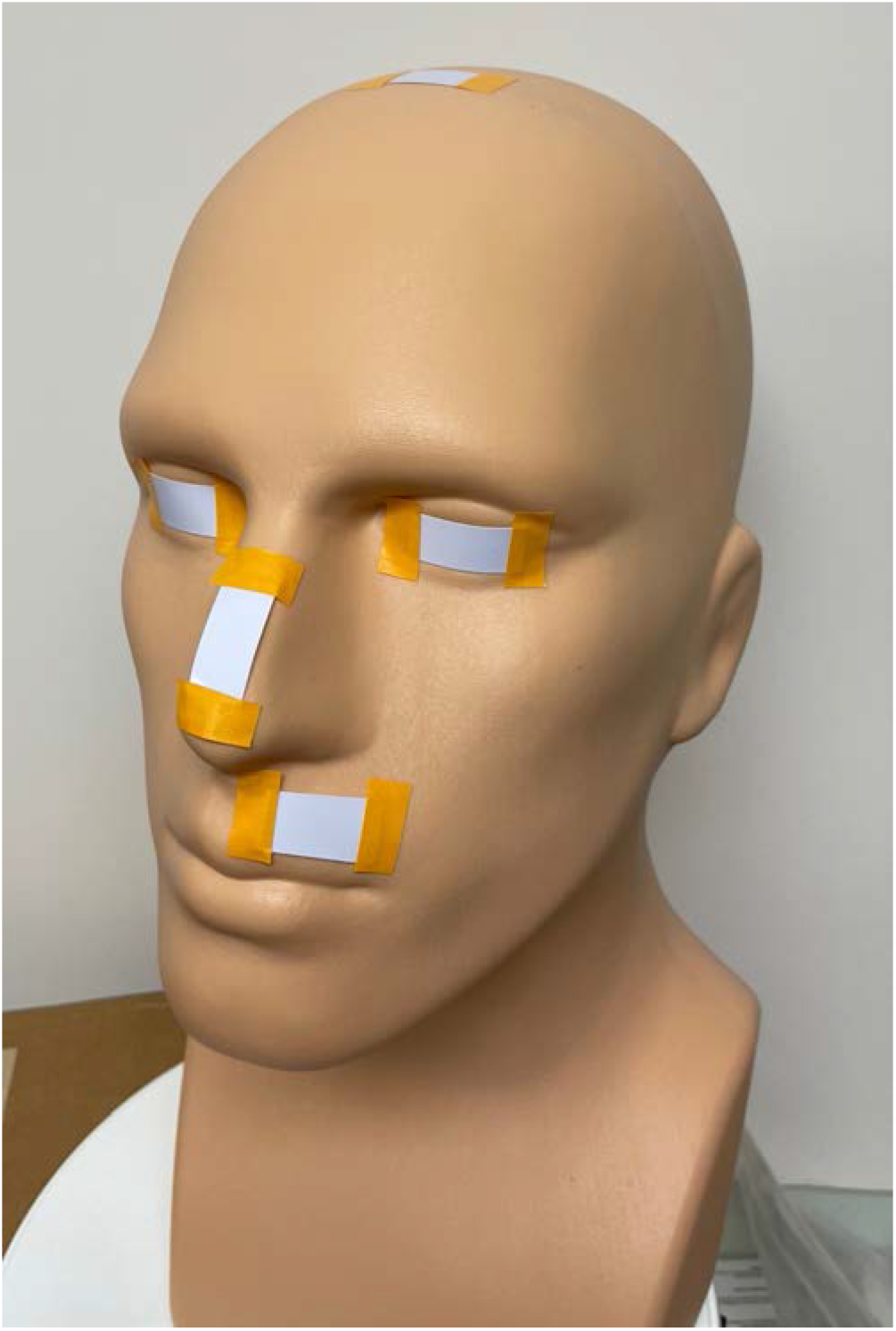
Film positions on the manikin. Films (light blue rectangles) were placed on the head, left and right eyes, nose and lip. Films were placed in the corresponding locations when the secondary manikin was used. Tape was used to affix the film to the manikin. Film on the head was placed on top of the hair when present, either from the addition of the wig or the in-built hair.

### Film analysis

We utilized an Epson Perfection V700 photo flatbed scanner (Epson, Suwa, NGN, Japan) for quantification of the color of each film(32). The scanner was operated in reflection mode and captured 48 bit RGB TIFF images with all color correction factors turned off. MATLAB (Mathworks, Natick, MA) was used for analysis. Only the green color channel was utilized for this study as it showed the best dose response. The color density (*CD*) of each pixel value was calculated with the equation:

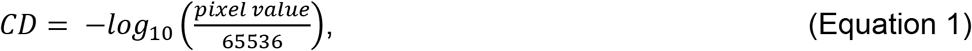

and the net color density calculated as:

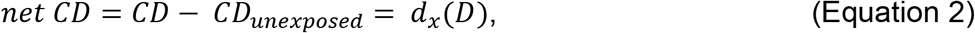

with the *CD_unexposed_* being the *CD* for an unexposed piece of film and *d_x_*(*D*) being the response for a given radiant exposure dose (*D*). Data for each exposure condition were matched to a fitting function with the form:

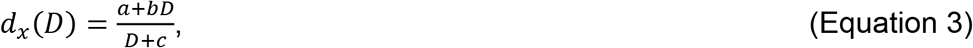

where *a*, *b*, and c are constants. The fitting function was optimized in MATLAB using the curve fitting tool to minimize the squared difference between the experimental data and the fit equation.

A calibration curve specific for the film dose response from exposure to the Lumenizer-300 fixture was established for use throughout testing. Calibration exposures were performed with an irradiance of 5 μW/cm^2^ measured using the UIT2400 meter. The radiant exposure doses for the calibration ranged from 135 μJ/cm^2^ to 5.67 mJ/cm^2^. The coefficients calculated to fit Equation 3 were *a*=0.00994 (95% confidence interval (CI) (−0.04648, 0.06637)), *b*=0.4424 (0.1989, 0.6859), and *c*=19.1 (5.786, 32.41), and the goodness of fit calculations produced an r-squared value of 0.9962.

The total radiant exposure dose received for each film was calculated. The radiant exposure dose was then divided by the exposure time to get the average irradiance upon the film during the 10-minute exposure. The average irradiance was then extrapolated to an 8-hour exposure to obtain the radiant exposure dose value reported for each film.

### Statistical analysis

A comprehensive statistical table was generated for the dose measured at each position using mean doses and standard deviations. Additionally, multivariate linear regression was performed (*R programing language* 4.0.3, Vienna, Austria) to model the responses of five dependent variables (doses in different facial locations) as function of six independent variables (predictors). The predictors were: sitting vs. standing (coded as sitting = 1, standing = 0), offset (0 meters, 0.5 meters, 1 meter), tilt (coded as no tilt = 0, 15 degrees forward = 1 and backwards = −1), manikin type (coded as primary manikin = 1, secondary manikin with in-built hair = 0), the presence of glasses applied to the primary manikin (coded as present = 1, absent = 0), and the presence of an afro wig applied to the primary manikin (coded as present = 1, absent = 0). The response variables were doses measured for each of the five film locations: the top of the head (head), the left cornea (left eye), the right cornea (right eye), the bridge of the nose (nose), and the superior lateral position of the upper lip (lip). Since the raw data for the response variables consisted of positive integer values, we used the Anscombe transformation to bring the data distribution closer to Gaussian to facilitate the linear regression analysis. The transformation is described as follows, where *D*(*i*) is the dose measured at location *i*, and *D_⊤_*(*i*) is the corresponding transformed dose, which was used as the response in multivariate regression:

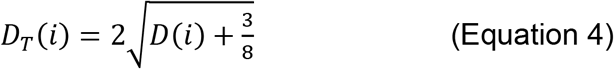

## Results

### Direct measurements of room irradiance

An assessment of the irradiance at the six manikin positions was performed using the optical power meter. Table 1 provides the results of the measured irradiance values and the extrapolated 8-hour doses. All assessment measurements maintained the experimental setup with the excimer lamps tilted at an outward angle of 45-degrees from the central axis of the lamp. The largest value for irradiance occurred at standing height (1.8 m) at an of offset 0.5 m with the meter aimed directly towards the fixture, which was recorded as the maximum directly measured dose. At this maximum, the directly measured 8-hour exposure dose was thus calculated to be 106.56 mJ/cm^2^.

**Table 1.** Measurements of the lamp using the optical power meter along with the corresponding calculated 8-hour doses. The maximum calculated 8-hour dose was chosen as the maximum directly measured dose (106.56 mJ/cm^2^ for this study).

| Measured Irradiance ($\mu\text{W}/\text{cm}^2$) | | | | |
| --- | --- | --- | --- | --- |
| Sensor configuration | Height | Offset |  |  |
|  |  | 0 m | 0.5 m | 1.0 m |
| Horizontal irradiance | 1.8 m | 3.100 | 3.100 | 0.300 |
|  | 1.3 m | 1.200 | 1.700 | 1.100 |
| Direct irradiance (meter directed at fixture) | 1.8 m | 3.100 | 3.700 | 0.550 |
|  | 1.3 m | 1.200 | 1.800 | 1.400 |
| Vertical irradiance | 1.8 m | 0.025 | 1.300 | 0.300 |
|  | 1.3 m | 0.018 | 0.260 | 0.530 |

| Calculated 8-hour dose (mJ/cm<sup>2</sup>) |  |  |  |  |
| --- | --- | --- | --- | --- |
| Sensor configuration | Height | Offset |  |  |
|  |  | 0 m | 0.5 m | 1.0 m |
| Horizontal irradiance | 1.8 m | 89.280 | 89.280 | 8.640 |
|  | 1.3 m | 34.560 | 48.960 | 31.680 |
| Direct irradiance (meter directed at fixture) | 1.8 m | 89.280 | **106.560** | 15.840 |
|  | 1.3 m | 34.560 | 51.840 | 40.320 |
| Vertical Irradiance | 1.8 m | 0.720 | 37.440 | 8.640 |
|  | 1.3 m | 0.518 | 7.488 | 15.264 |

### Measurement of radiant exposure using manikins

Color density analysis of OrthoChromic OC-1 film exposure from the 222nm KrCl lamp was performed as described above. Extrapolated 8-hour doses for all the measurements are provided in the Supplemental Materials Table S1. The measured doses ranged from 0 mJ/cm^2^ to 183 mJ/cm^2^. Aggregate doses for extrapolated 8-hour exposures (in mJ/cm^2^) for all manikin conditions are plotted for the various positions in Figure 4. The values for the data plotted in Figure 4 are included in the Supplemental Materials as Table S2. The highest average radiant exposure doses for the sitting and standing position were at an offset of 0.5-meters away from the source and on top of the head, with doses of 59.25 mJ/cm^2^ (sd= 10.88) and 150.4 mJ/cm^2^ (sd= 24.5) respectively. Eye doses remained close to zero in every sitting position, with the highest average doses at a 1-meter offset for both the left eye and right eye of 0.92 mJ/cm^2^ (sd= 2.1) and 1.42 mJ/cm^2^ (sd= 2.23), respectively. Interestingly, the highest exposures to the left and right eye in the standing position took place at an offset of 0.5-meters with doses of 5.17 mJ/cm^2^ (sd= 9.1) and 7.33 mJ/cm^2^ (sd= 11.6), respectively.

**Figure 4.**
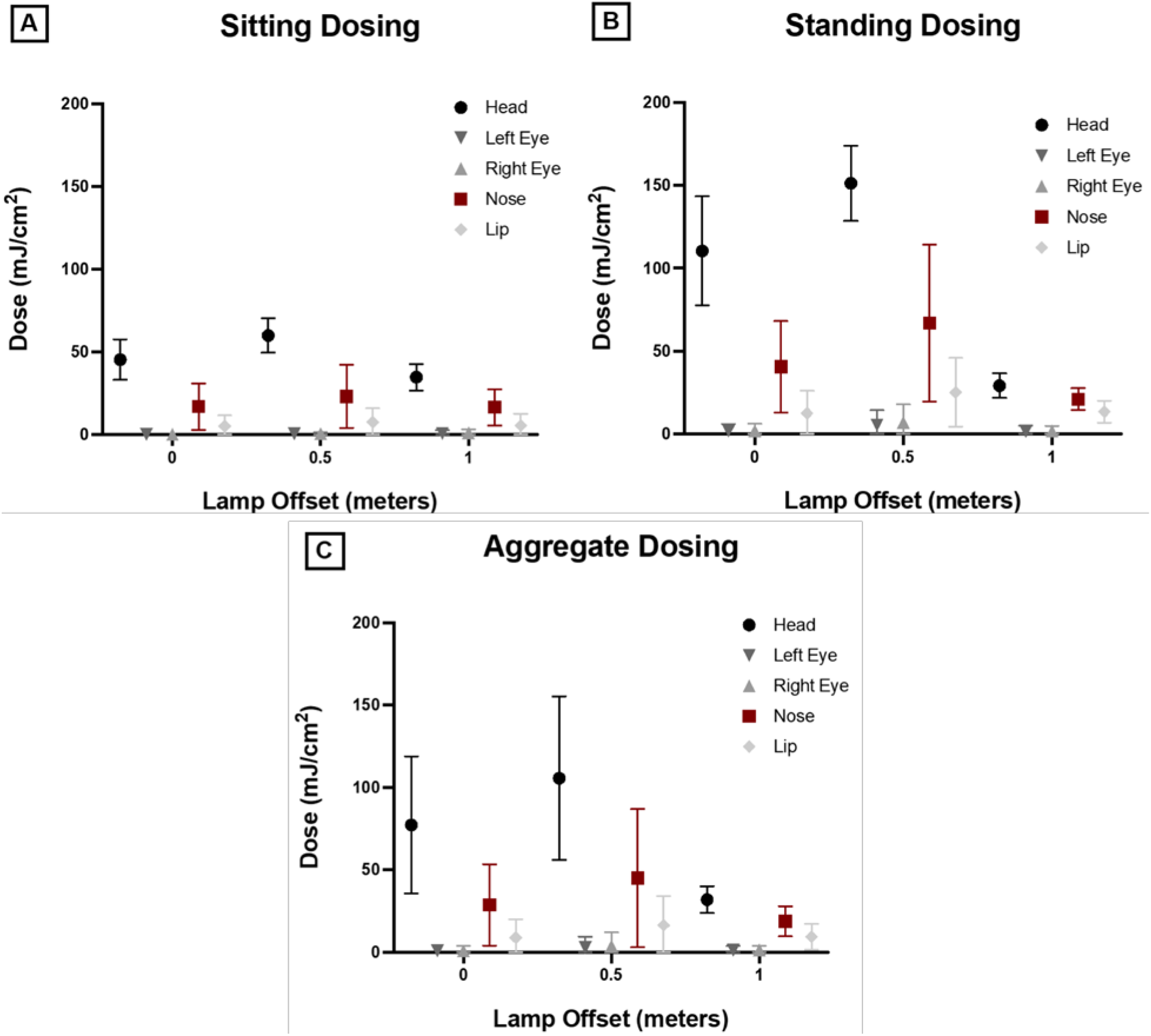
Summary of aggregated dosing values. Average extrapolated 8-hour film dosing for manikin positions of sitting(A), standing(B), and aggregate of sitting and standing (C). Mean doses with representative standard deviation error bars are plotted. The results are also included as Table S2 within the Supplemental Materials.

### Predictor dependent dosing effects: a multivariate linear regression

A multivariate linear regression was used to predict the effects of the six predictor variables on the dosing to the five anatomical locations. Results of the regression are provided in Table 2. Sitting was shown to be the only consistent significant predictor for dosing in all five film locations with the head showing the largest effect from sitting (the largest absolute value of the slope coefficient, β) and the right eye showing the smallest effect from sitting (β=-5.1 standard error (SE)=0.97, p= 1.93E-06 and β=-1.14 SE=0.41, p=6.81E-03) respectively. Tilting of the manikin 15 degrees backwards showed a significant effect to all film locations except for the top of the head, with the largest effect seen on the nose (β=-3.35 SE=0.47, p=1.27E-9). Additionally, offset showed a significant effect to dosing of the head (β=-4.81 SE=1.19, p=1.51E-04) and the lip (β=-2.98 SE=0.48, p=3.61 E-08). Finally, the presence or absence of hair or a wig showed significant effects to the right eye, nose, and lip, while glasses showed a marginally significant effect only to the right eye.

**Table 2.** Multivariate linear regression results. Significant predictor p-values are bolded.

| Film Location | Predictor | $\beta$ Coeff. | Std.err. | p-value |
| --- | --- | --- | --- | --- |
| Head | Intercept | 19.81 | 1.24 | 3.77E-24 |
|  | Sitting | -5.10 | 0.97 | **1.93E-06** |
|  | Offset | -4.81 | 1.19 | **1.51E-04** |
|  | Tilt | 0.13 | 0.60 | 8.26E-01 |
|  | Primary manikin | 1.33 | 1.38 | 3.39E-01 |
|  | Glasses | 0.65 | 1.38 | 6.37E-01 |
|  | Primary manikin with wig | -0.36 | 1.38 | 7.95E-01 |
| Left Eye | Intercept | 2.38 | 0.50 | 1.07E-05 |
|  | Sitting | -1.16 | 0.39 | **4.21E-03** |
|  | Offset | 0.31 | 0.48 | 5.26E-01 |
|  | Tilt | -1.12 | 0.24 | **1.48E-05** |
|  | Primary manikin | 0.99 | 0.55 | 7.79E-02 |
|  | Glasses | -1.06 | 0.55 | 6.08E-02 |
|  | Primary manikin with wig | -1.10 | 0.55 | 5.14E-02 |
| Right Eye | Intercept | 2.24 | 0.52 | 5.36E-05 |
|  | Sitting | -1.14 | 0.41 | **6.81E-03** |
|  | Offset | 0.50 | 0.50 | 3.15E-01 |
|  | Tilt | -1.25 | 0.25 | **4.24E-06** |
|  | Primary manikin | 1.49 | 0.57 | **1.20E-02** |
|  | Glasses | -1.34 | 0.57 | **2.27E-02** |
|  | Primary manikin with wig | -1.35 | 0.57 | **2.16E-02** |
| Nose | Intercept | 7.27 | 0.99 | 3.66E-10 |
|  | Sitting | -3.94 | 0.77 | **3.20E-06** |
|  | Offset | 0.18 | 0.95 | 8.49E-01 |
|  | Tilt | -3.35 | 0.47 | **1.27E-09** |
|  | Primary manikin | 6.62 | 1.09 | **7.77E-08** |
|  | Glasses | 0.36 | 1.09 | 7.45E-01 |
|  | Primary manikin with wig | -3.86 | 1.09 | **7.65E-04** |
| Lip | Intercept | 4.64 | 0.61 | 1.27E-10 |
|  | Sitting | -2.98 | 0.48 | **3.61E-08** |
|  | Offset | 1.31 | 0.58 | **2.85E-02** |
|  | Tilt | -3.25 | 0.29 | **1.02E-16** |
|  | Primary manikin | 3.02 | 0.67 | **3.00E-05** |
|  | Glasses | 0.23 | 0.67 | 7.33E-01 |
|  | Primary manikin with wig | -1.68 | 0.67 | **1.49E-02** |

## Discussion

This study measured the UV dose received from a 222 nm far-UVC excimer lamp fixture ceiling-mounted to directly expose an occupied area. This setup allowed for measurement of the predicted 8-hour exposure dose on a manikin at five anatomical positions including the top of the head, the left and right eye, the bridge of the nose, and the superior lateral portion of the lip. The results indicated that regardless of manikin position the highest dose was received upon the top of the head, with the next highest dose upon the bridge of the nose, followed by the superior lateral position of the lip, and finally, the eyes.

As with installations of upper room UV, any installation of far-UVC within an occupied space will need to operate within the TLVs or ELs. Upon review of the results of this work, some of the 8-hour doses from the current exposure setup exceeded the 23 mJ/cm^2^ ICNIRP EL for 222 nm radiation, particularly those for the top of the head in the standing position. We will note that these measurements did not account for any movement around the room through time weighted averaging, which is standard practice when applying the TLV or EL (47). Therefore, an assumption that an occupant would perhaps be standing in the exposure area for only 1 hour out of 8 would reduce the expected dose proportionally. As with upper room UV installations, a careful assessment of the entire installation taking into account factors such as expected positioning of occupants within a space would be needed for any far-UVC installations for direct human exposure.

The most sensitive organ to UVC damage is the eye, and accordingly the TLVs and ELs have been set to prevent eye irritation (33, 34). Notable within the conditions tested in this work is the minimal exposure dose upon the eye. As shown in Figure 4 and Supplemental Table S2, when the dose is aggregated from all the measured positions the dose received by the eyes is well below the 23 mJ/cm^2^ ICNIRP EL for 222 nm radiation. However, the constant rotation of the manikin head during the test exposure presents a possible weakness of this study in measuring dose to the eye and other facial features. The constant rotation was included to capture multiple exposure angles to try to represent realistic exposure scenarios; however, the rotation also resulted in the face pointing away from the source for half of the exposure time. It may be appropriate to multiply the exposure doses for the eye and face by a factor of two to account for this. Notably, one exposure condition did result in an 8-hour dose to the eye of greater than the 23 mJcm^2^ ICNIRP EL for 222 nm: the standing height of 1.8 m at an offset of 0.5 m with the manikin in a tilted back position. However, accounting for time weighted averaging, since it would be highly unlikely someone would stay in this position staring into the fixture for 8-hours, would reduce these exposure doses below the daily EL.

A useful metric for assessing eye hazard is to look at the dose received by the eyes as a percentage of the maximum directly measured dose. Measurements for direct exposure dose in this work suggested a maximum 8-hour dose of 106.56 mJ/cm^2^. The average dose received by the eyes was maximum when standing 0.5 m offset from the lamp, with an average eye dose of 6.25 mJ/cm^2^ (left eye 5.17 mJ/cm^2^, right eye 7.33 mJ/cm^2^) across all of the conditions tested; this suggests doses to the eye of only 5.8% of the directly measured dose. This result is in good agreement with the measurements by Urbach for orbital exposure from the sun, which reported between 0-10% radiation to the orbit compared to the maximum exposure dose received (48). Similarly, Sydenham *et al.* reported the eye receiving approximately 22% of the solar dose to the chest (49). Overall, it appears that the anatomical features which protect human eyes from solar radiation will provide similar protection for ceiling-mounted installations of far-UVC.

The 2022 publication of the ACGIH TLVs included an increase to the ultraviolet radiation limits in much of the UVC (36). The new recommendations separate the TLVs for eye exposure from those for skin exposure. The new TLV for skin exposure at 222 nm, about 479 mJ/cm^2^, is approximately three times the TLV for eye at 222 nm, which is about 160 mJ/cm^2^. The results of this work showed an average dose to the eye of only 5.8% of the maximum direct dose, which suggests that the eye dose should not approach the TLV as long as the direct measurements remain within the TLV, using the respective TLVs. Since the ICNIRP EL does not make a distinction between eye and skin limits, the dose to the eyes when exposing with a direct exposure at the EL would likewise only be a fraction of the allowable dose.

The position with the highest dose to the eyes was at standing height and 0.5 m offset. This maximum is likely due to both the emission pattern of the lamp, with radiation emitting at an angle away from the fixture, and the protective anatomical features of the brow being less effective with radiation from a lower angle. Also important to note when reviewing these results is that while the manikin has many anatomically correct features, the manikin omitted any protective features of the eyelid. In reality, changes in the eyelid position and periodic blinking would further reduce the actual dose received upon the eye.

The multivariate linear regression analysis performed in this work permitted an evaluation of the impact of each of the predictor variables tested upon the expected dose at each anatomical position on the manikin. As expected, changing the distance from the source between the sitting and standing positions was significant for predicting dose to all film locations. While offset of the manikin had a significant effect on dose to the head and lip, it was not a significant factor for other film locations; however, this could be caused by the small values for doses recorded for locations such as the eye limiting the measurement range. The tilt of the manikin backwards was associated with higher exposures to the eyes, nose, and lip, since these features would be more directly exposed with this head position. Comparing the two manikin types yielded some significant differences in the right eye, nose, and lip. These results suggest that anatomical differences, such as brow ridge shape, can ultimately effect dose received by an individual. The addition of glasses to the manikin did show a decrease in dose received by the eyes, but the effect was only significant in the left eye; again, the small maximum dose received by the eye limited the overall dose range that could be analyzed so large decreases were not possible. Finally, the addition of the afro wig to the manikin decreased the received dose in many of the film locations that were protected by the hair. It is notable that the film for the top of the head was placed on top of the hair during exposures with the wig, so this dose is not indicative of what would be received at the scalp, which should be significantly reduced by the presence of hair.

While this study encompassed various manikin positions and features, we stress that these results should not be used as a statement regarding the hazard of operation of this particular fixture. Instead, each installation of far-UVC should be installed and commissioned to operate within the TLVs or ELs within the unique space they are placed.

Overall this study provides the first analysis of possible exposure doses which might be experienced using an indoor far-UVC installation. The use of manikin head phantoms to simulate the natural protection of the eyes by the brow and other anatomical features makes this a good model for expected eye dose. Importantly, the results indicate that the eyes only receive a small fraction of directly measured dose, calculated at 5.8% in this study, and this can be useful for hazard assessment with future far-UVC installations.

## Acknowledgements

Funding for this work was provided by NIH grant 5R42AI125006, NASA grant 80NSSC22K0211, Columbia University Urban Tech Award: Technology Innovations for Better Urban Living, and the Shostack Foundation.

## References

1. Kowalski, W., Ultraviolet Germicidal Irradiation Handbook: UVGI for Air and Surface Disinfection. Springer. 2009, New York: Springer.

2. Buonanno, M., B. Ponnaiya, D. Welch, M. Stanislauskas, G. Randers-Pehrson, L. Smilenov, F.D. Lowy, D.M. Owens, and D.J. Brenner, Germicidal Efficacy and Mammalian Skin Safety of 222-nm UV Light. Radiat Res, 2017. 187(4): p. 483–491.

3. Buonanno, M., G. Randers-Pehrson, A.W. Bigelow, S. Trivedi, F.D. Lowy, H.M. Spotnitz, S.M. Hammer, and D.J. Brenner, 207-nm UV light - a promising tool for safe low-cost reduction of surgical site infections. I: in vitro studies. PloS one, 2013. 8(10): p. e76968–e76968.

4. Buonanno, M., D. Welch, I. Shuryak, and D.J. Brenner, Far-UVC light (222 nm) efficiently and safely inactivates airborne human coronaviruses. Scientific reports, 2020. 10(1): p. 10285–8.

5. Welch, D., M. Buonanno, V. Grilj, I. Shuryak, C. Crickmore, A.W. Bigelow, G. Randers-Pehrson, G.W. Johnson, and D.J. Brenner, Far-UVC light: A new tool to control the spread of airborne-mediated microbial diseases. Scientific Reports, 2018. 8(1): p. 2752.

6. Welch, D., M. Buonanno, I. Shuryak, G. Randers-Pehrson, H.M. Spotnitz, and D.J. Brenner, Effect of far ultraviolet light emitted from an optical diffuser on methicillin-resistant Staphylococcus aureus in vitro. PloS one, 2018. 13(8): p. e0202275–e0202275.

7. Ponnaiya, B., M. Buonanno, D. Welch, I. Shuryak, G. Randers-Pehrson, and D.J. Brenner, Far-UVC light prevents MRSA infection of superficial wounds in vivo. PloS one, 2018. 13(2): p. e0192053–e0192053.

8. Welch, D., M. Aquino de Muro, M. Buonanno, and D.J. Brenner, Wavelength-dependent DNA Photodamage in a 3-D human Skin Model over the Far-UVC and Germicidal UVC Wavelength Ranges from 215 to 255 nm. Photochemistry and Photobiology, 2022.

9. Narita, K., K. Asano, K. Naito, H. Ohashi, M. Sasaki, Y. Morimoto, T. Igarashi, and A. Nakane, Ultraviolet C light with wavelength of 222 nm inactivates a wide spectrum of microbial pathogens. The Journal of hospital infection, 2020. 105(3): p. 459–467.

10. Taylor, W., E. Camilleri, D.L. Craft, G. Korza, M.R. Granados, J. Peterson, R. Szczpaniak, S.K. Weller, R. Moeller, T. Douki, W.W.K. Mok, and P. Setlow, DNA Damage Kills Bacterial Spores and Cells Exposed to 222-Nanometer UV Radiation. Applied and environmental microbiology, 2020. 86(8).

11. Kang, J.-W. and D.-H. Kang, The Synergistic Bactericidal Mechanism of Simultaneous Treatment with a 222-Nanometer Krypton-Chlorine Excilamp and a 254-Nanometer Low-Pressure Mercury Lamp. Applied and environmental microbiology, 2019. 85(1).

12. Sosnin, E., S. Avdeev, E. Kuznetzova, and L. Lavrent’eva, A Bactericidal Barrier-Discharge KrBr Excilamp. Instruments and experimental techniques (New York), 2005. 48(5): p. 663–666.

13. Buchan, A.G., L. Yang, and K.D. Atkinson, Predicting airborne coronavirus inactivation by far-UVC in populated rooms using a high-fidelity coupled radiation-CFD model. Scientific Reports, 2020. 10(1): p. 19659.

14. Eadie, E., I.M. Barnard, S.H. Ibbotson, and K. Wood, Extreme exposure to filtered far-UVC: A case study. Photochemistry and Photobiology, 2021. 97(3): p. 527–531.

15. Eadie, E., P. O’Mahoney, L. Finlayson, I.R.M. Barnard, S.H. Ibbotson, and K. Wood, Computer Modeling Indicates Dramatically Less DNA Damage from Far-UVC Krypton Chloride Lamps (222 nm) than from Sunlight Exposure. Photochemistry and Photobiology, 2021.

16. Yamano, N., M. Kunisada, S. Kaidzu, K. Sugihara, A. Nishiaki-Sawada, H. Ohashi, A. Yoshioka, T. Igarashi, A. Ohira, M. Tanito, and C. Nishigori, Long-term Effects of 222-nm ultraviolet radiation C Sterilizing Lamps on Mice Susceptible to Ultraviolet Radiation. Photochem Photobiol, 2020. 96(4): p. 853–862.

17. Yamano, N., M. Kunisada, A. Nishiaki-Sawada, H. Ohashi, T. Igarashi, and C. Nishigori, Evaluation of Acute Reactions on Mouse Skin Irradiated with 222 and 235 nm UV-C. Photochemistry and Photobiology, 2021. 97(4): p. 770–777.

18. Yam, J.C. and A.K. Kwok, Ultraviolet light and ocular diseases. Int Ophthalmol, 2014. 34(2): p. 383–400.

19. Sengillo, J.D., A.L. Kunkler, C. Medert, B. Fowler, M. Shoji, N. Pirakitikulr, N. Patel, N.A. Yannuzzi, A.J. Verkade, D. Miller, D.H. Sliney, J.M. Parel, and G. Amescua, UV-Photokeratitis Associated with Germicidal Lamps Purchased during the COVID-19 Pandemic. Ocul Immunol Inflamm, 2021. 29(1): p. 76–80.

20. Rubeshkumar, P.C., M. Ponnaiah, D. Anandhi, and D. John, Association between exposure to artificial sources of ultraviolet radiation and ocular diseases: a systematic review protocol. JBI Evid Synth, 2020. 18(8): p. 1766–1773.

21. Boulton, M., M. Rózanowska, and B. Rózanowski, Retinal photodamage. J Photochem Photobiol B, 2001. 64(2-3): p. 144–61.

22. Cullen, A.P., Photokeratitis and other phototoxic effects on the cornea and conjunctiva. Int J Toxicol, 2002. 21(6): p. 455–64.

23. Roberts, J.E., Ocular phototoxicity. J Photochem Photobiol B, 2001. 64(2-3): p. 136–43.

24. Sliney, D.H., Exposure geometry and spectral environment determine photobiological effects on the human eye. Photochem Photobiol, 2005. 81(3): p. 483–9.

25. McDevitt, J.J., S.N. Rudnick, and L.J. Radonovich, Aerosol susceptibility of influenza virus to UV-C light. Appl Environ Microbiol, 2012. 78(6): p. 1666–9.

26. Yin, R., T. Dai, P. Avci, A.E. Jorge, W.C. de Melo, D. Vecchio, Y.Y. Huang, A. Gupta, and M.R. Hamblin, Light based anti-infectives: ultraviolet C irradiation, photodynamic therapy, blue light, and beyond. Curr Opin Pharmacol, 2013. 13(5): p. 731–62.

27. Glasfeld, A., Biochemistry: The Chemical Reactions of Living Cells, 2nd Edition (David E. Metzler). Journal of Chemical Education, 2004. 81 (5): p. 646.

28. Goldfarb, A.R. and L.J. Saidel, Ultraviolet absorption spectra of proteins. Science, 1951. 114(2954): p. 156–7.

29. Buonanno, M., M. Stanislauskas, B. Ponnaiya, A.W. Bigelow, G. Randers-Pehrson, Y. Xu, I. Shuryak, L. Smilenov, D.M. Owens, and D.J. Brenner, 207-nm UV Light-A Promising Tool for Safe Low-Cost Reduction of Surgical Site Infections. II: In-Vivo Safety Studies. PLoS One, 2016. 11(6): p. e0138418.

30. Doughty, M.J. and M.L. Zaman, Human corneal thickness and its impact on intraocular pressure measures: a review and meta-analysis approach. Surv Ophthalmol, 2000. 44(5): p. 367–408.

31. Kolozsvári, L., A. Nógrádi, B. Hopp, and Z. Bor, UV absorbance of the human cornea in the 240-to 400-nm range. Invest Ophthalmol Vis Sci, 2002. 43(7): p. 2165–8.

32. Welch, D. and D.J. Brenner, Improved Ultraviolet Radiation Film Dosimetry Using OrthoChromic OC-1 Film(dagger). Photochem Photobiol, 2021. 97(3): p. 498–504.

33. Threshold Limit Values and Biological Exposure Indices. American Conference of Governmental Industrial Hygienists., 2012.

34. Guidelines on limits of exposure to ultraviolet radiation of wavelengths between 180 nm and 400 nm (incoherent optical radiation). Health Phys, 2004. 87(2): p. 171–86.

35. American Conference of Governmental Industrial Hygienists, 2021 Threshold Limit Values and Biological Exposure Indices. 2021, Cincinnati, Ohio: American Conference of Governmental Industrial Hygienists.

36. American Conference of Governmental Industrial Hygienists, 2022 Threshold Limit Values and Biological Exposure Indices. 2022, Cincinnati, Ohio: American Conference of Governmental Industrial Hygienists.

37. Rosenthal, F.S., C. Phoon, A.E. Bakalian, and H.R. Taylor, The ocular dose of ultraviolet radiation to outdoor workers. Invest Ophthalmol Vis Sci, 1988. 29(4): p. 649–56.

38. Linde, K., C.Y. Wright, T. Kapwata, and J.L. du Plessis, Low Use of Ocular Sun Protection among Agricultural Workers in South Africa: Need for Further Research. Photochem Photobiol, 2021. 97(2): p. 453–455.

39. Deng, Y., C. Zhang, Y. Zheng, R. Li, H. Hua, Y. Lu, N. Gurram, R. Chen, N. OuYang, S. Zhang, Y. Liu, and L. Hu, Effect of Protective Measures on Eye Exposure to Solar Ultraviolet Radiation. Photochem Photobiol, 2021. 97(1): p. 205–212.

40. Downs, N.J., A.V. Parisi, P.W. Schouten, D.P. Igoe, and G. De Castro-Maqueda, The Simulated Ocular and Whole-Body Distribution of Natural Sunlight to Kiteboarders: A High-Risk Case of UVR Exposure for Athletes Utilizing Water Surfaces in Sport. Photochem Photobiol, 2020. 96(4): p. 926–935.

41. Herlihy, E., P.H. Gies, C.R. Roy, and M. Jones, PERSONAL DOSIMETRY OF SOLAR UV RADIATION FOR DIFFERENT OUTDOOR ACTIVITIES. Photochemistry and Photobiology, 1994. 60(3): p. 288–294.

42. Kimlin, M.G., A.V. Parisi, and J.C.F. Wong, Quantification of personal solar UV exposure of outdoor workers, indoor workers and adolescents at two locations in Southeast Queensland. Photodermatology, photoimmunology & photomedicine, 1998. 14(1): p. 7–11.

43. Peters, C., S. Kalia, P. Demers, A.-M. Nicol, and M. Koehoorn, 0211 Solar ultraviolet radiation (UVR) exposure levels and sun protection behaviours in outdoor workers in British Columbia, Canada. Occupational and environmental medicine (London, England), 2014. 71 (Suppl 1): p. A27–A28.

44. Diffey, B.L., C.J. Gibson, R. Haylock, and A.F. McKinlay, Outdoor ultraviolet exposure of children and adolescents. British Journal of Dermatology, 1996. 134(6): p. 1030–1034.

45. Parisi, A.V., M.G. Kimlin, R. Lester, and D. Turnbull, Lower body anatomical distribution of solar ultraviolet radiation on the human form in standing and sitting postures. J Photochem Photobiol B, 2003. 69(1): p. 1–6.

46. Vital and Health Statistics. 2021, NATIONAL CENTER FOR HEALTH STATISTICS: Center for Disease Control and Prevention.

47. First, M.W., R.A. Weker, S. Yasui, and E.A. Nardell, Monitoring Human Exposures to Upper-Room Germicidal Ultraviolet Irradiation. Journal of Occupational and Environmental Hygiene, 2005. 2(5): p. 285–292.

48. Urbach, F., Geographic distribution of skin cancer. Journal of surgical oncology, 1971. 3(3): p. 219–234.

49. Sydenham, M.M., M.J. Collins, and L.W. Hirst, Measurement of ultraviolet radiation at the surface of the eye. Investigative ophthalmology & visual science, 1997. 38(8): p. 1485–1492.

