## Supplemental Materials for "Ocular and Facial Far-UVC Doses from Ceiling-Mounted 222 nm Far-UVC Fixtures"

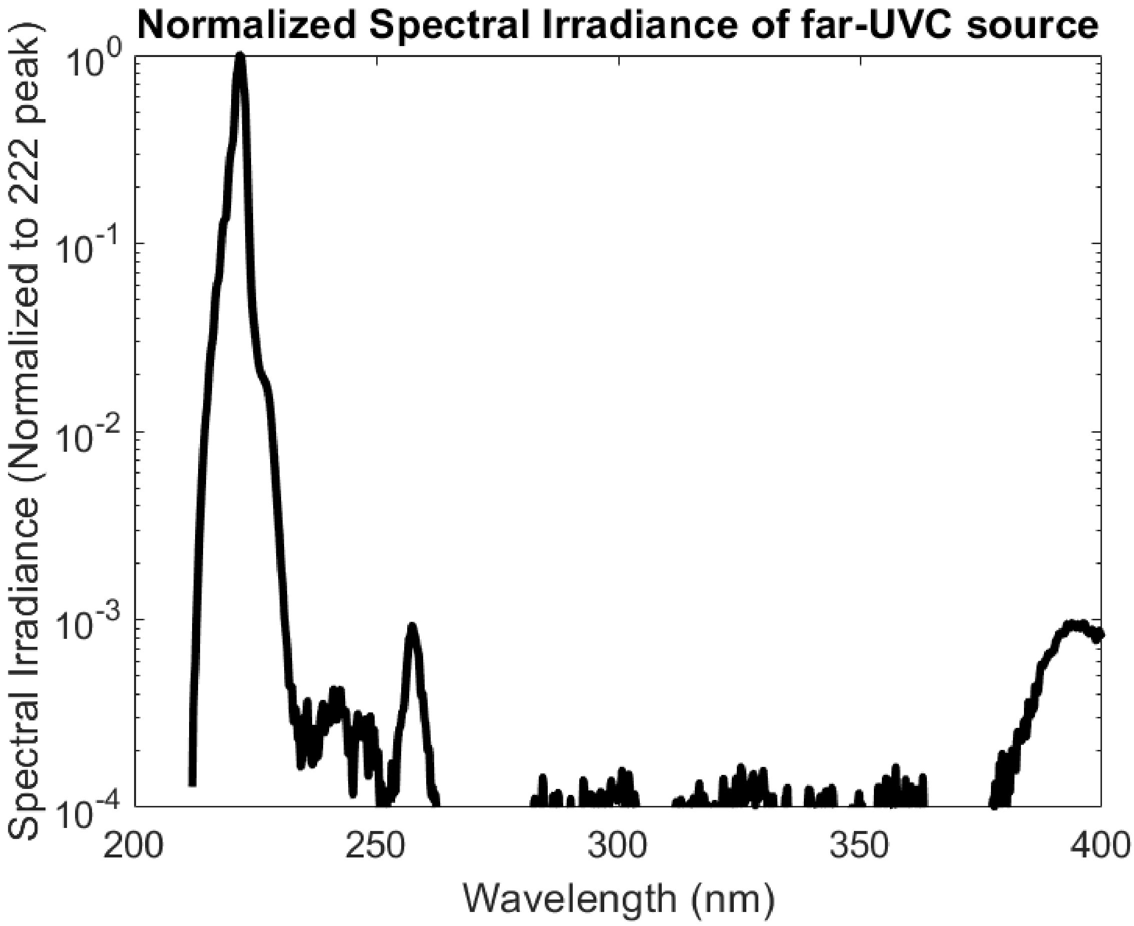


**Figure S1**. The emission spectrum for the far-UVC lamp used throughout this study. The spectrum is normalized to the 222 nm peak. The minimal output outside of the peak is indicative of filtering of the characteristic KrCl excimer emission spectrum.

**Table S1.** The complete dataset acquired in this study. Measurements of the 8-hour extrapolated dose to the head, eyes, nose, and lip are given for each configuration tested. All doses are expressed in mJ/cm$^2$. The last column relates the eye dose as a percentage of the maximum directly measured dose.

| Sitting/ Standing | Offset | Tilt | Manikin | Glasses | Afro Wig | Head | Left Eye | Right Eye | Nose | Lip | Eye Average | Max Dose Directly Measured | Eye % of Max Directly Measured |
|---|---|---|---|---|---|---|---|---|---|---|---|---|---|
| sitting | 0 | 0 | Primary | No | No | 48 | 0 | 0 | 24 | 10 | 0 | 106.56 | 0% |
| sitting | 0 | Back | Primary | No | No | 46 | 0 | 0 | 30 | 19 | 0 | 106.56 | 0% |
| sitting | 0 | Front | Primary | No | No | 44 | 0 | 0 | 16 | 0 | 0 | 106.56 | 0% |
| sitting | 0.5 | 0 | Primary | No | No | 70 | 0 | 0 | 30 | 14 | 0 | 106.56 | 0% |
| sitting | 0.5 | Back | Primary | No | No | 64 | 1 | 2 | 53 | 21 | 1.5 | 106.56 | 1% |
| sitting | 0.5 | Front | Primary | No | No | 64 | 0 | 0 | 16 | 0 | 0 | 106.56 | 0% |
| sitting | 1 | 0 | Primary | No | No | 35 | 0 | 2 | 24 | 10 | 1 | 106.56 | 1% |
| sitting | 1 | Back | Primary | No | No | 37 | 2 | 4 | 28 | 17 | 3 | 106.56 | 3% |
| sitting | 1 | Front | Primary | No | No | 40 | 0 | 0 | 14 | 0 | 0 | 106.56 | 0% |
| standing | 0 | 0 | Primary | No | No | 85 | 6 | 2 | 64 | 14 | 4 | 106.56 | 4% |
| standing | 0 | Back | Primary | No | No | 113 | 16 | 15 | 60 | 25 | 15.5 | 106.56 | 15% |
| standing | 0 | Front | Primary | No | No | 113 | 1 | 1 | 36 | 0 | 1 | 106.56 | 1% |
| standing | 0.5 | 0 | Primary | No | No | 165 | 0 | 5 | 90 | 25 | 2.5 | 106.56 | 2% |
| standing | 0.5 | Back | Primary | No | No | 145 | 25 | 38 | 122 | 50 | 31.5 | 106.56 | 30% |
| standing | 0.5 | Front | Primary | No | No | 153 | 3 | 2 | 46 | 12 | 2.5 | 106.56 | 2% |
| standing | 1 | 0 | Primary | No | No | 25 | 0 | 0 | 17 | 6 | 0 | 106.56 | 0% |
| standing | 1 | Back | Primary | No | No | 25 | 9 | 11 | 20 | 20 | 10 | 106.56 | 9% |
| standing | 1 | Front | Primary | No | No | 20 | 0 | 1 | 23 | 16 | 0.5 | 106.56 | 0% |
| sitting | 0 | 0 | Secondary | No | No | 30 | 0 | 0 | 0 | 0 | 0 | 106.56 | 0% |
| sitting | 0 | Back | Secondary | No | No | 30 | 0 | 0 | 3 | 0 | 0 | 106.56 | 0% |
| sitting | 0 | Front | Secondary | No | No | 39 | 0 | 0 | 0 | 0 | 0 | 106.56 | 0% |
| sitting | 0.5 | 0 | Secondary | No | No | 52 | 0 | 0 | 0 | 0 | 0 | 106.56 | 0% |
| sitting | 0.5 | Back | Secondary | No | No | 56 | 2 | 2 | 19 | 5 | 2 | 106.56 | 2% |
| sitting | 0.5 | Front | Secondary | No | No | 60 | 0 | 0 | 0 | 0 | 0 | 106.56 | 0% |
| sitting | 1 | 0 | Secondary | No | No | 29 | 0 | 1 | 9 | 0 | 0.5 | 106.56 | 0% |

| Sitting/ Standing | Offset | Tilt | Manikin | Glasses | Afro Wig | Head | Left Eye | Right Eye | Nose | Lip | Eye Average | Max Dose Directly Measured | Eye % of Max Directly Measured |
|---|---|---|---|---|---|---|---|---|---|---|---|---|---|
| sitting | 1 | Back | Secondary | No | No | 35 | 7 | 7 | 11 | 11 | 7 | 106.56 | 7% |
| sitting | 1 | Front | Secondary | No | No | 48 | 0 | 0 | 2 | 0 | 0 | 106.56 | 0% |
| standing | 0 | 0 | Secondary | No | No | 74 | 0 | 0 | 0 | 0 | 0 | 106.56 | 0% |
| standing | 0 | Back | Secondary | No | No | 53 | 0 | 0 | 33 | 13 | 0 | 106.56 | 0% |
| standing | 0 | Front | Secondary | No | No | 91 | 0 | 0 | 0 | 0 | 0 | 106.56 | 0% |
| standing | 0.5 | 0 | Secondary | No | No | 120 | 0 | 0 | 17 | 10 | 0 | 106.56 | 0% |
| standing | 0.5 | Back | Secondary | No | No | 136 | 2 | 3 | 56 | 44 | 2.5 | 106.56 | 2% |
| standing | 0.5 | Front | Secondary | No | No | 97 | 0 | 0 | 0 | 0 | 0 | 106.56 | 0% |
| standing | 1 | 0 | Secondary | No | No | 47 | 0 | 1 | 28 | 7 | 0.5 | 106.56 | 0% |
| standing | 1 | Back | Secondary | No | No | 26 | 10 | 2 | 16 | 21 | 6 | 106.56 | 6% |
| standing | 1 | Front | Secondary | No | No | 31 | 0 | 0 | 10 | 2 | 0 | 106.56 | 0% |
| sitting | 0 | 0 | Primary | Yes | No | 50 | 0 | 0 | 28 | 13 | 0 | 106.56 | 0% |
| sitting | 0 | Back | Primary | Yes | No | 35 | 0 | 0 | 31 | 12 | 0 | 106.56 | 0% |
| sitting | 0 | Front | Primary | Yes | No | 51 | 0 | 0 | 9 | 0 | 0 | 106.56 | 0% |
| sitting | 0.5 | 0 | Primary | Yes | No | 68 | 0 | 0 | 26 | 6 | 0 | 106.56 | 0% |
| sitting | 0.5 | Back | Primary | Yes | No | 64 | 2 | 3 | 56 | 21 | 2.5 | 106.56 | 2% |
| sitting | 0.5 | Front | Primary | Yes | No | 60 | 0 | 0 | 7 | 1 | 0 | 106.56 | 0% |
| sitting | 1 | 0 | Primary | Yes | No | 47 | 0 | 0 | 25 | 5 | 0 | 106.56 | 0% |
| sitting | 1 | Back | Primary | Yes | No | 35 | 0 | 0 | 40 | 15 | 0 | 106.56 | 0% |
| sitting | 1 | Front | Primary | Yes | No | 40 | 0 | 0 | 19 | 2 | 0 | 106.56 | 0% |
| standing | 0 | 0 | Primary | Yes | No | 98 | 0 | 0 | 61 | 25 | 0 | 106.56 | 0% |
| standing | 0 | Back | Primary | Yes | No | 119 | 3 | 4 | 74 | 36 | 3.5 | 106.56 | 3% |
| standing | 0 | Front | Primary | Yes | No | 99 | 0 | 0 | 30 | 0 | 0 | 106.56 | 0% |
| standing | 0.5 | 0 | Primary | Yes | No | 183 | 0 | 8 | 99 | 25 | 4 | 106.56 | 4% |
| standing | 0.5 | Back | Primary | Yes | No | 159 | 10 | 10 | 138 | 57 | 10 | 106.56 | 9% |
| standing | 0.5 | Front | Primary | Yes | No | 168 | 0 | 0 | 41 | 5 | 0 | 106.56 | 0% |
| standing | 1 | 0 | Primary | Yes | No | 37 | 3 | 0 | 29 | 26 | 1.5 | 106.56 | 1% |
| standing | 1 | Back | Primary | Yes | No | 36 | 0 | 0 | 27 | 20 | 0 | 106.56 | 0% |
| standing | 1 | Front | Primary | Yes | No | 36 | 0 | 0 | 29 | 11 | 0 | 106.56 | 0% |
| sitting | 0 | 0 | Primary | No | yes | 36 | 0 | 0 | 0 | 0 | 0 | 106.56 | 0% |
| sitting | 0 | Back | Primary | No | yes | 45 | 0 | 0 | 29 | 11 | 0 | 106.56 | 0% |

| Sitting/ Standing | Offset | Tilt | Manikin | Glasses | Afro Wig | Head | Left Eye | Right Eye | Nose | Lip | Eye Average | Max Dose Directly Measured | Eye % of Max Directly Measured |
|---|---|---|---|---|---|---|---|---|---|---|---|---|---|
| sitting | 0 | Front | Primary | No | yes | 27 | 0 | 0 | 0 | 0 | 0 | 106.56 | 0% |
| sitting | 0.5 | 0 | Primary | No | yes | 30 | 0 | 0 | 10 | 0 | 0 | 106.56 | 0% |
| sitting | 0.5 | Back | Primary | No | yes | 54 | 0 | 2 | 42 | 16 | 1 | 106.56 | 1% |
| sitting | 0.5 | Front | Primary | No | yes | 69 | 0 | 0 | 0 | 0 | 0 | 106.56 | 0% |
| sitting | 1 | 0 | Primary | No | yes | 29 | 0 | 0 | 17 | 3 | 0 | 106.56 | 0% |
| sitting | 1 | Back | Primary | No | yes | 24 | 2 | 3 | 28 | 20 | 2.5 | 106.56 | 2% |
| sitting | 1 | Front | Primary | No | yes | 41 | 0 | 0 | 2 | 0 | 0 | 106.56 | 0% |
| standing | 0 | 0 | Primary | No | yes | 96 | 0 | 0 | 24 | 2 | 0 | 106.56 | 0% |
| standing | 0 | Back | Primary | No | yes | 166 | 0 | 0 | 66 | 20 | 0 | 106.56 | 0% |
| standing | 0 | Front | Primary | No | yes | 111 | 0 | 0 | 0 | 0 | 0 | 106.56 | 0% |
| standing | 0.5 | 0 | Primary | No | yes | 175 | 0 | 0 | 47 | 25 | 0 | 106.56 | 0% |
| standing | 0.5 | Back | Primary | No | yes | 139 | 22 | 22 | 107 | 44 | 22 | 106.56 | 21% |
| standing | 0.5 | Front | Primary | No | yes | 165 | 0 | 0 | 0 | 0 | 0 | 106.56 | 0% |
| standing | 1 | 0 | Primary | No | yes | 30 | 0 | 0 | 26 | 14 | 0 | 106.56 | 0% |
| standing | 1 | Back | Primary | No | yes | 24 | 1 | 3 | 16 | 14 | 2 | 106.56 | 2% |
| standing | 1 | Front | Primary | No | yes | 23 | 0 | 0 | 21 | 8 | 0 | 106.56 | 0% |

**Table S2.** Summary of exposure data plotted in Figure 4. Values are expressed as the extrapolated 8-hour dose in $mJ/cm^2$.

| | Head | | | | Left Eye | | | Right Eye | | | Nose | | | Lip | | |
|---|---|---|---|---|---|---|---|---|---|---|---|---|---|---|---|---|
| | Offset | Mean | SD | N | Mean | SD | N | Mean | SD | N | Mean | SD | N | Mean | SD | N |
| **Sitting** | 0 | 40.08 | 8.38 | 12 | 0.00 | 0.00 | 12 | 0.00 | 0.00 | 12 | 14.2 | 13.5 | 12 | 5.42 | 7.03 | 12 |
| | 0.5 | 59.25 | 10.88 | 12 | 0.42 | 0.8 | 12 | 0.75 | 1.14 | 12 | 21.6 | 20.1 | 12 | 7.00 | 8.57 | 12 |
| | 1 | 36.67 | 7.15 | 12 | 0.92 | 2.1 | 12 | 1.42 | 2.23 | 12 | 18.3 | 11.4 | 12 | 6.92 | 7.38 | 12 |
| | Offset | Mean | SD | N | Mean | SD | N | Mean | SD | N | Mean | SD | N | Mean | SD | N |
| **Standing** | 0 | 101.5 | 27.6 | 12 | 2.17 | 4.7 | 12 | 1.83 | 4.32 | 12 | 37.3 | 27.6 | 12 | 11.3 | 12.8 | 12 |
| | 0.5 | 150.4 | 24.5 | 12 | 5.17 | 9.1 | 12 | 7.33 | 11.6 | 12 | 63.6 | 46.9 | 12 | 24.8 | 20.03 | 12 |
| | 1 | 30.00 | 7.76 | 12 | 1.92 | 3.7 | 12 | 1.50 | 3.15 | 12 | 21.8 | 6.19 | 12 | 13.8 | 7.20 | 12 |
| | Offset | Mean | SD | N | Mean | SD | N | Mean | SD | N | Mean | SD | N | Mean | SD | N |
| **Aggregate** | 0 | 70.79 | 37.2 | 24 | 1.08 | 3.5 | 24 | 0.92 | 3.13 | 24 | 25.8 | 24.3 | 24 | 8.33 | 10.5 | 24 |
| | 0.5 | 104.8 | 50.1 | 24 | 2.79 | 6.7 | 24 | 4.04 | 8.75 | 24 | 42.6 | 41.3 | 24 | 15.9 | 17.6 | 24 |
| | 1 | 33.33 | 8.05 | 24 | 1.42 | 2.9 | 24 | 1.46 | 2.67 | 24 | 20.04 | 9.15 | 24 | 10.3 | 7.94 | 24 |
